# Draft genome sequences of four lactic acid bacteria from fermented chicken meat unveil biosynthetic gene clusters for antimicrobial compounds

**DOI:** 10.1101/2025.01.18.633471

**Authors:** Mahabub Alam, Tamanna Hassan, Tanvir Ahmad Nizami, Lipi Akter, Tofazzal Md Rakib

## Abstract

Lactic acid bacteria play a crucial role in fermented food production and serve as important sources of antimicrobial peptides. This study reports four lactic acid bacteria strains, isolated from fermented chicken meat, which harbor biosynthetic gene clusters encoding antimicrobial compounds. These strains are classified within the genera *Pediococcus* and *Lactiplantibacillus*.

## Announcement

Fermented foods have long carried probiotics, prebiotics, and postbiotics. Lactic acid bacteria (LAB) enhance food safety, inhibit spoilage, and support gut health. LAB, from the phylum *Firmicutes*, include *Lactobacillus, Pediococcus, Leuconostoc, Lactococcus*, and *Enterococcus* (1, 2). With rising antibiotic resistance, bacteriocins are gaining attention as potential alternatives (3). Genome mining and high-throughput screening of biosynthetic gene clusters present a promising pathway for discovering novel bacteriocins, addressing resistance challenges, and advancing sustainable food preservation (4, 5). Three broiler chickens from a live bird market (22.336218 N, 91.830187 E) in Chattogram, Bangladesh, were slaughtered under standard procedures. Breast meat (100 g) was stored anaerobically at 4°C for three days to propagate probiotic bacteria. One gram of meat was mixed with 5 mL MRS broth (Oxoid, Hampshire, England) and incubated anaerobically at 37°C for 48 hours, promoting LAB growth. Single colonies were isolated on MRS agar, purified, and confirmed by genus-specific 16S rRNA amplification. Genomic DNA was extracted from 1.5 mL of 24-hour cultured broth using a AllPrep Bacterial DNA/RNA/Protein Kit (Qiagen, Hilden, Germany), and purity was assessed with a Nanodrop One (Thermo Fisher Scientific, MA, USA). DNA libraries were prepared using the Nextera XT DNA Library Preparation Kit (Illumina, CA, USA). Genome sequencing was conducted on an Illumina NextSeq2000 platform at the Poultry Research and Training Center, Chattogram Veterinary and Animal Sciences University, Bangladesh, generating 2×150 bp paired-end reads. Raw data quality, including read quality, GC content, and adapter contamination, was assessed using FastQC v0.12.1 (6). Trim Galore v0.6.5dev (7) was employed to trim low-quality bases and adapter sequences, followed by normalization of read coverage with bbnorm v39.08 (8) to reduce redundancy. De novo genome assembly was performed using Unicycler v0.4.8 (9), with subsequent polishing using Pilon v1.24 (10) to enhance accuracy. Genome assembly quality was assessed using QUAST v5.2.0 (11), while Samtools v1.13 (12) was used for alignment file manipulation and indexing. The de novo genome assembly, conducted through the BV-BRC (13) comprising above tools, yielded coverage ranging from 65.4× to 145.6×, with raw reads between 1,452,975 and 1,662,521. The assembled genomes were annotated using NCBI PGAP v6.9 (14) and PATRIC (15) with the in-built RASTtk pipeline to identify the species, with transport proteins identified via TCDB (16), drug targets predicted using DrugBank v6.0 (17), and additional analyses performed for CRISPR arrays and prophages using CRISPRCasFinder (18), and Integrative and Conjugative Elements using ICEFinder (19). All software parameters were set to default unless specified otherwise. Detailed genomic characteristics are provided in Table 1.

**Table 1:**
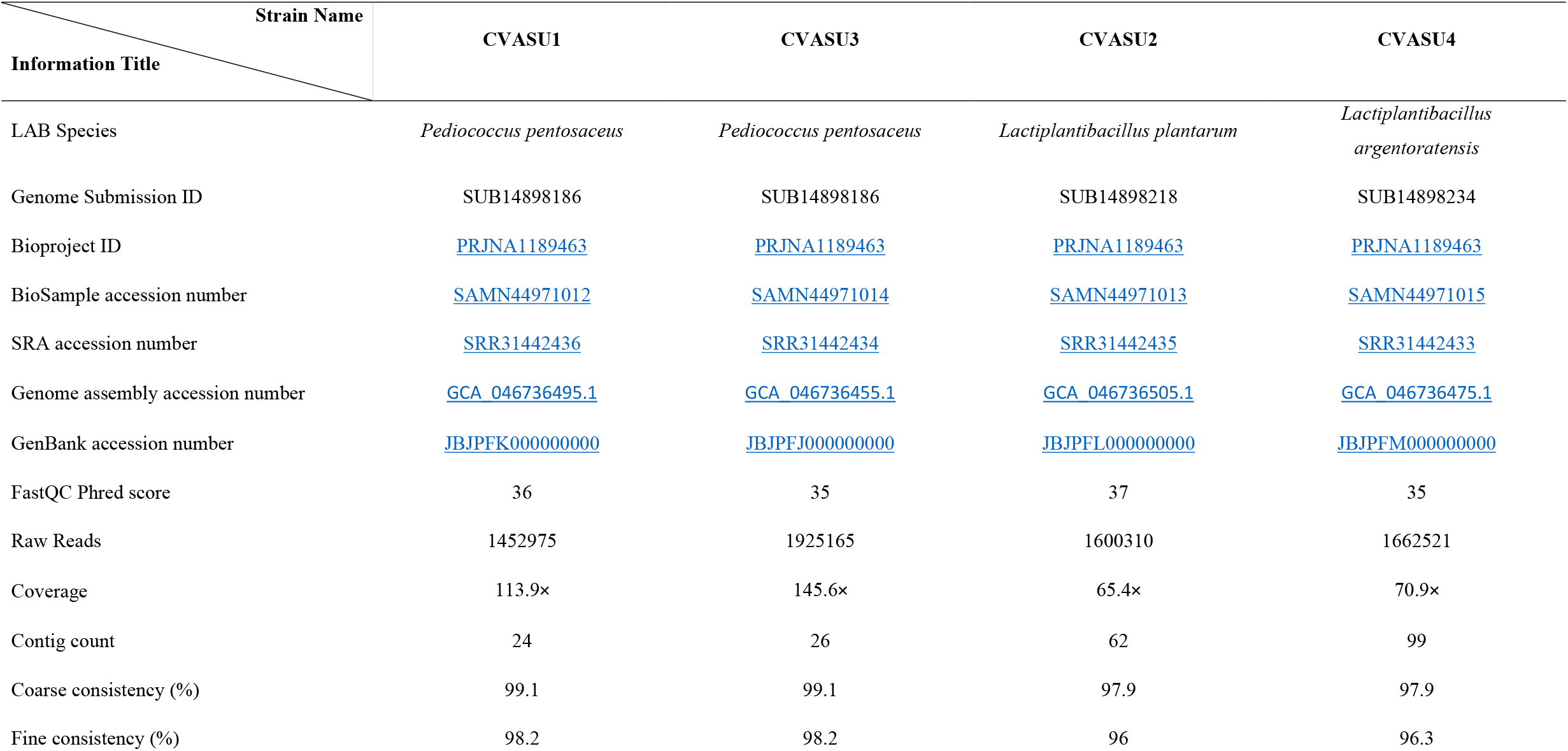
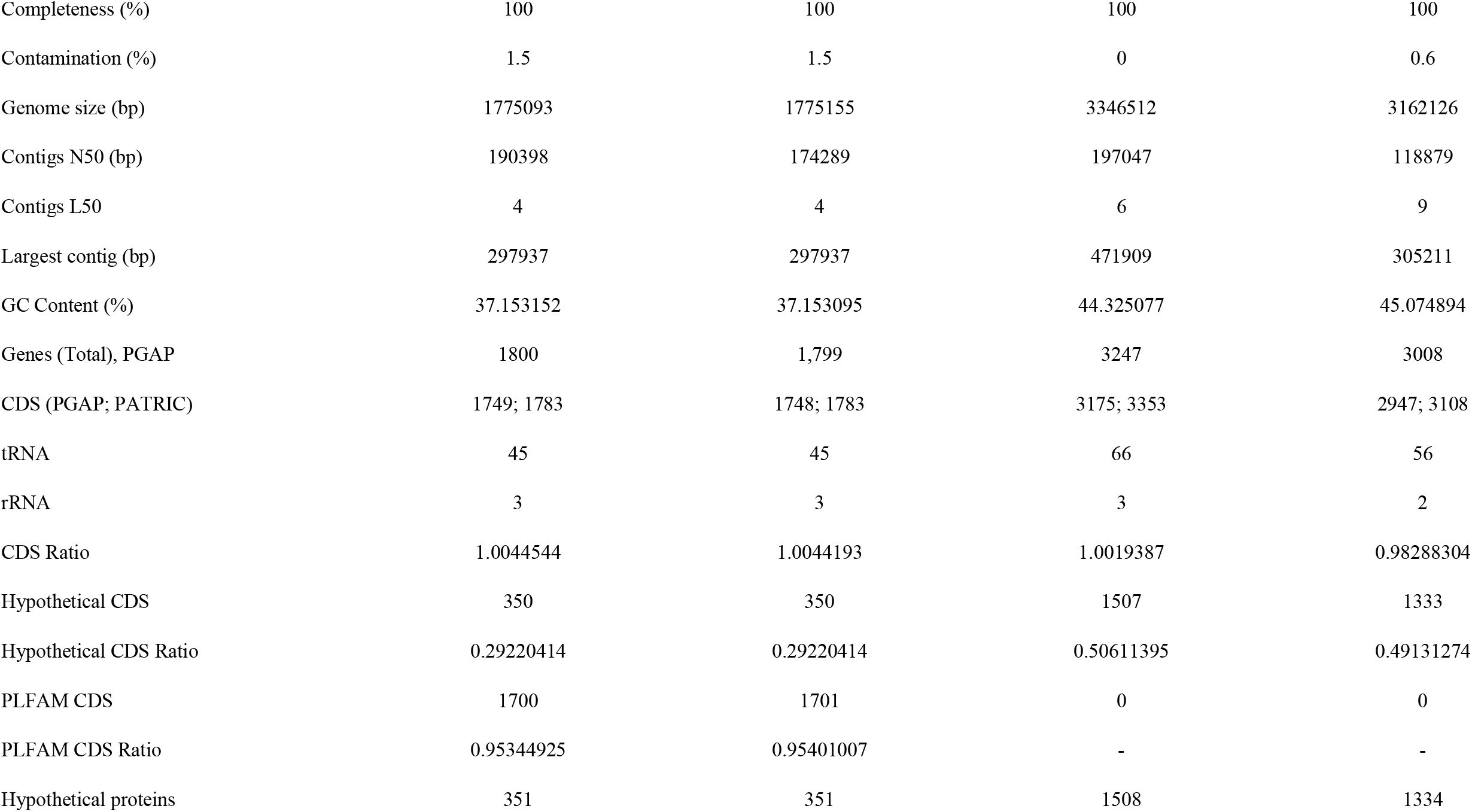
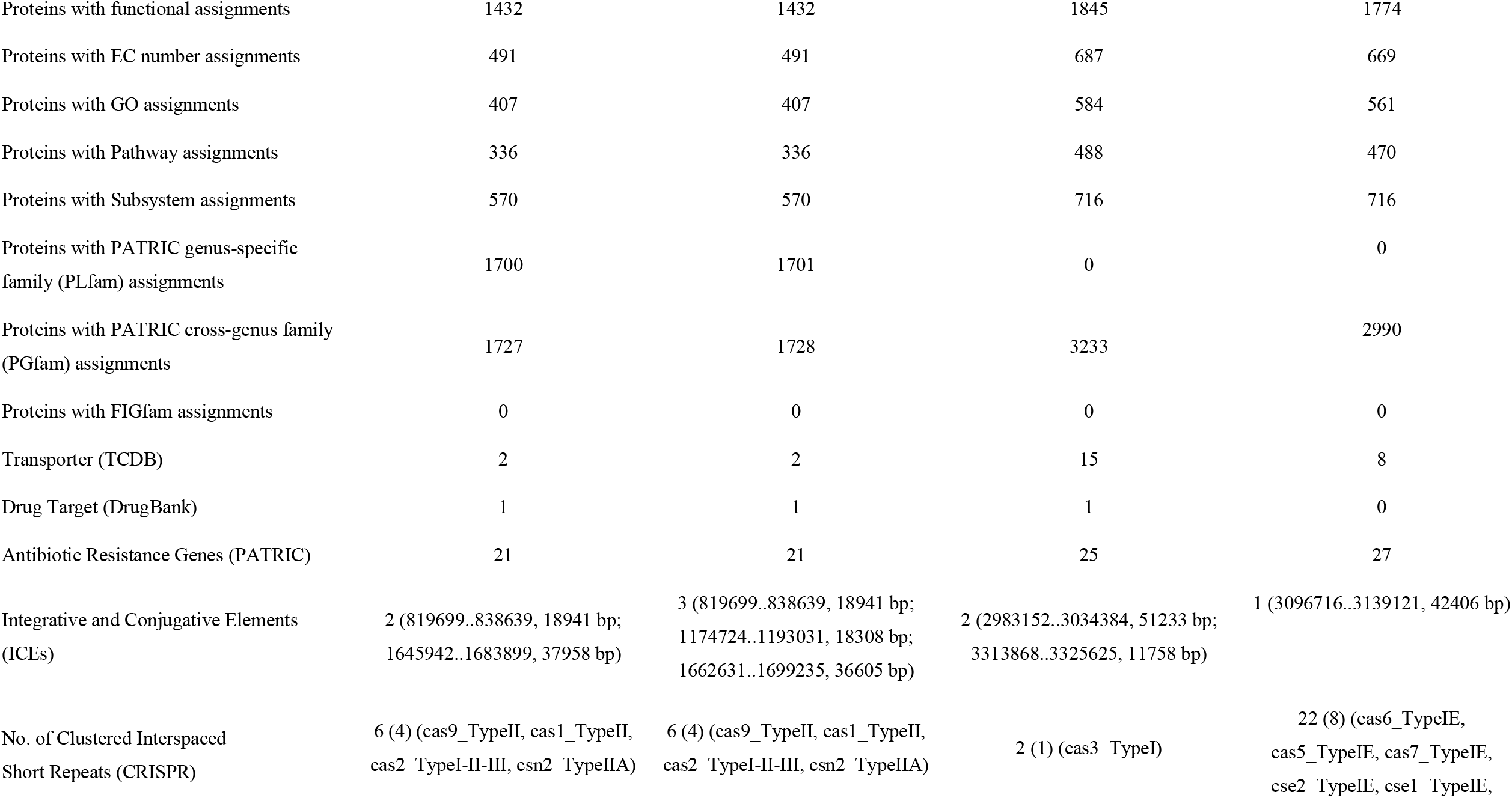
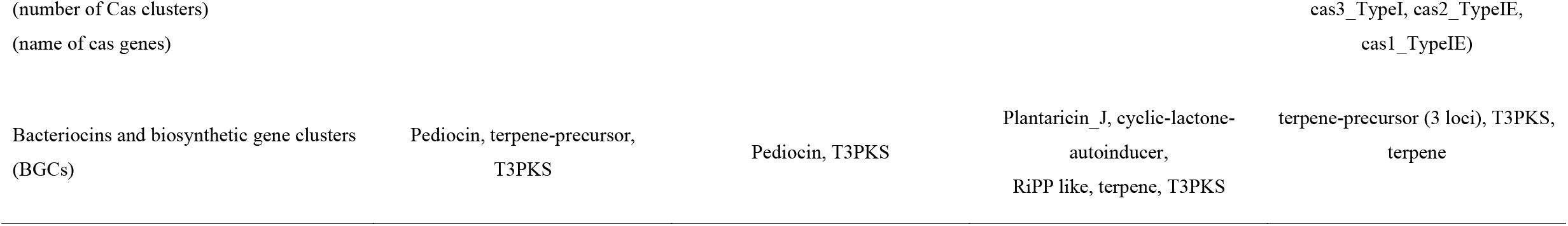
Genomic features of the sequenced four LAB strains isolated from fermented chicken meat in Chattogram, Bangladesh.

The genomes of four LAB isolates were mined for biosynthetic gene clusters (BGCs) associated with antimicrobial production using the software antiSMASH v7.0 (20) and BAGEL4 (21). The genomes were found to contain multiple BGCs encoding ribosomally synthesized and posttranslationally modified peptides (RiPP like compounds), Cyclic Lactone Autoinducer, Type III Polyketide Synthase (T3PKS), Terpene Precursors, Terpene, and Class IIa (Pediocin) and IIb (Plantaricin_J) bacteriocins. The isolates or their purified bacteriocins may be used in the future for food preservation and commercial probiotics to manage gut health and related diseases.

## Data availability statement

The accession numbers for the genomes of the four LAB strains are shown in Table 1. All genomes are available under BioProject accession number PRJNA1189463.

## Acknowledgments

The genome sequencing was supported by Chattogram Veterinary and Animal Sciences University WGS establishment project.

## References

Zhao J, Zhao J, Zang J, Peng C, Li Z, Zhang P. 2024. Isolation, identification, and evaluation of lactic acid bacteria with probiotic potential from traditional fermented sour meat. Frontiers in Microbiology 15:1421285

Fernandes A, Jobby R. 2022. Bacteriocins from lactic acid bacteria and their potential clinical applications. Applied Biochemistry and Biotechnology 194:4377–4399

Cleveland J, Montville TJ, Nes IF, Chikindas ML. 2001. Bacteriocins: safe, natural antimicrobials for food preservation. International journal of food microbiology 71:1–20

Gerst M, Yousef A. 2018. Modified microassay for the isolation of antimicrobial-producing, spore-forming and nonspore-forming bacteria. Journal of applied microbiology 124:1401–1410

Zhou H, Fang J, Tian Y, Lu XY. 2014. Mechanisms of nisin resistance in Gram-positive bacteria. Annals of microbiology 64:413–420

Andrews S. 2010. FastQC: A quality control tool for high throughput sequence data. URL https://wwwbioinformaticsbabrahamacuk/projects/fastqc/.

Krueger F. 2019. A wrapper tool around Cutadapt and FastQC to consistently apply quality and adapter trimming to FastQ files, with some extra functionality for MspI-digested RRBS-type (Reduced Representation Bisufite-Seq) libraries. URL https://wwwbioinformaticsbabrahamacuk/projects/trim_galore/.

Bushnell B. 2017. BBNorm: Normalize coverage by down-sampling reads over high-depth areas of a genome, to result in a flat coverage distribution. Department of Energy, Joint Genome Institute, CA, USA. URL https://jgidoegov/data-and-tools/software-tools/bbtools/bb-tools-user-guide/bbnorm-guide/.

Wick RR, Judd LM, Gorrie CL, Holt KE. 2017. Unicycler: resolving bacterial genome assemblies from short and long sequencing reads. PLoS computational biology 13:e1005595

Walker BJ, Abeel T, Shea T, Priest M, Abouelliel A, Sakthikumar S, Cuomo CA, Zeng Q, Wortman J, Young SK. 2014. Pilon: an integrated tool for comprehensive microbial variant detection and genome assembly improvement. PloS one 9:e112963

Gurevich A, Saveliev V, Vyahhi N, Tesler G. 2013. QUAST: quality assessment tool for genome assemblies. Bioinformatics 29:1072–1075

Danecek P, Bonfield JK, Liddle J, Marshall J, Ohan V, Pollard MO, Whitwham A, Keane T, McCarthy SA, Davies RM. 2021. Twelve years of SAMtools and BCFtools. Gigascience 10:giab008

Olson RD, Assaf R, Brettin T, Conrad N, Cucinell C, Davis JJ, Dempsey DM, Dickerman A, Dietrich EM, Kenyon RW. 2023. Introducing the bacterial and viral bioinformatics resource center (BV-BRC): a resource combining PATRIC, IRD and ViPR. Nucleic acids research 51:D678–D689

Tatusova T, DiCuccio M, Badretdin A, Chetvernin V, Nawrocki EP, Zaslavsky L, Lomsadze A, Pruitt KD, Borodovsky M, Ostell J. 2016. NCBI prokaryotic genome annotation pipeline. Nucleic acids research 44:6614–6624

Wattam AR, Davis JJ, Assaf R, Boisvert S, Brettin T, Bun C, Conrad N, Dietrich EM, Disz T, Gabbard JL. 2017. Improvements to PATRIC, the all-bacterial bioinformatics database and analysis resource center. Nucleic acids research 45:D535–D542

Saier Jr MH, Reddy VS, Moreno-Hagelsieb G, Hendargo KJ, Zhang Y, Iddamsetty V, Lam KJK, Tian N, Russum S, Wang J. 2021. The transporter classification database (TCDB): 2021 update. Nucleic acids research 49:D461–D467

Knox C, Wilson M, Klinger CM, Franklin M, Oler E, Wilson A, Pon A, Cox J, Chin NE, Strawbridge SA. 2024. DrugBank 6.0: the DrugBank knowledgebase for 2024. Nucleic acids research 52:D1265–D1275

Couvin D, Bernheim A, Toffano-Nioche C, Touchon M, Michalik J, Néron B, Rocha EP, Vergnaud G, Gautheret D, Pourcel C. 2018. CRISPRCasFinder, an update of CRISRFinder, includes a portable version, enhanced performance and integrates search for Cas proteins. Nucleic acids research 46:W246–W251

Gonçalves OS, de Assis JCS, Santana MF. 2022. Breaking the ICE: an easy workflow for identifying and analyzing integrative and conjugative elements in bacterial genomes. Functional & Integrative Genomics 22:1139–1145

Blin K, Shaw S, Medema MH, Weber T. 2024. The antiSMASH database version 4: additional genomes and BGCs, new sequence-based searches and more. Nucleic acids research 52:D586–D589

van Heel AJ, de Jong A, Song C, Viel JH, Kok J, Kuipers OP. 2018. BAGEL4: a user-friendly web server to thoroughly mine RiPPs and bacteriocins. Nucleic acids research 46:W278–W281

